# Mutant huntingtin protein alters the response of microglial cells to inflammatory stimuli

**DOI:** 10.1101/550913

**Authors:** David W. Donley, Ryan Nelson, Jason P. Gigley, Jonathan H. Fox

## Abstract

Huntington’s disease (HD) is a progressive neurodegenerative disease that affects the striatum and cerebral cortex. It is caused by a dominant CAG trinucleotide expansion in exon 1 of the *HTT* gene. Mutant huntingtin protein (mHtt) is expressed in neurons and immune cells. HD patients demonstrate altered blood cytokine profiles and altered responses of peripheral immune cells to inflammatory stimuli. However, the effects of mHtt on microglial immune responses are not fully understood. Herein we discuss the current understanding of how mHtt alters microglial inflammatory responses. Using lentivirus, we expressed the N171 N-terminal fragment of wild-type or mhtt containing 18 and 82 glutamine repeats in cultured EOC-20 microglial cells. We then measured responses to lipopolysaccharide or interleukin-6. Mutant huntingtin-expressing microglial cells produced less interleukin-6 and nitric oxide in response to lipopolysaccharide stimulation than wild-type huntingtin-expressing cells. However, mHtt-expressing microglia stimulated with interleukin-6 produced more nitric oxide than wild-type cells. Mutant huntingtin-expressing cells had higher basal NF-κB and further elevations of NF-κB after interleukin-6 but not lipopolysaccharide stimulation. Thus we demonstrate the potential of mHtt to dampen responses to lipopolysaccharide but potentiate responses to interleukin-6. This work adds to the emerging understanding that mHtt alters not only baseline status of cells but may also result in altered immune responses dependent on the nature of the inflammatory stimuli. We also present our perspective that in human HD the extent of inflammation may depend, in part, on altered responses to varied inflammatory stimuli including environmental factors such as infection.

## Inflammation in Huntington’s disease

Huntington’s disease is caused by a genetically dominant CAG repeat expansion in the huntingtin gene (*HTT*) that results in expression of polyglutamine-expanded mutant huntingtin protein (mHtt) in neurons and microglial cells (Cisbani and Cicchetti, 2012). Neuroinflammation, marked by microglial activation, is an early feature of Huntington’s disease (HD) (Sapp et al., 2001, Tai et al., 2007). Further, neuroinflammation may potentiate neurodegenerative processes and promote HD progression. HD-positive individuals exhibit increased systemic inflammation, marked by elevated interleukin 6 and other cytokines, that begins years before clinical onset (Björkqvist et al., 2008). However, the relationship between peripheral and brain inflammation in HD is poorly understood. Further, the extent to which inflammation in HD results from intrinsic effect of mHtt versus altered responses to environmental stimuli is unclear.

## Monocytic cells in Huntington’s disease

Microglial and monocytes are innate immune cells that may contribute to HD pathogenesis. Microglia are monocyte-like CNS-resident cells and express many of the same surface receptors and markers as peripheral monocytes (Greter et al., 2015). Peripheral monocytes are cellular precursors of tissue-resident macrophages, dendritic cells, monocyte-derived suppressor cells and infiltrating microglial-like cells in the CNS (Greter et al., 2015). Both microglial cells and monocytes can be activated by immune mediators as well as directly by microbial molecules.

Mutant huntingtin is expressed in both microglia and monocytes in HD patients (Weiss et al., 2012). In the absence of external inflammatory stimulation, mHtt promotes cell-autonomous activation of primary microglial cells (Crotti et al., 2014). Immune cells from pre-manifest HD patients and mouse models demonstrate a pro-inflammatory phenotype as illustrated by increased levels of several cytokines in blood, including interleukin-6 (IL-6) (Crotti et al., 2014, Björkqvist et al., 2008). Mutant huntingtin expression in human monocytes is associated with increased production of cytokines interleukin-6 (IL-6), IL-1β, and TNF-α (Träger et al., 2014). Ĳnterleukin-6 induces microglial proliferation and is associated with increased microglia proliferation in response to mHtt-expressing neurons (Kraft et al., 2012). Furthermore, monocytes isolated from HD patients and mice had reduced chemotactic responses to ATP and complement protein C5a further demonstrating that mHtt exerts a modifying effect on these cells (Kwan et al., 2012). Together, these data indicate that HD monocytes and microglia have changes in basal activation state and that mHtt may alter how monocyte-derived cells respond to innate immune stimuli and/or sensitivity to cytokine and chemokine stimulation.

Macrophages can be polarized in their response to be either “M1” or “M2” cells (Mills et al., 2000). In HD, expression of CCR2 and production of the pro-inflammatory cytokine IL-12 are used to identify M1 macrophages while CX3CR1 expression and immunosuppressive IL-10 production are used to identify M2 macrophages (Di Pardo et al., 2013). Interestingly, HD patient monocytes are initially more M1 polarized as evidenced by increased percentages of CCR2+ and IL-12+ macrophages before disease onset, with a transition to M2 macrophages that express CX3CR1 and IL-10 later in the disease course (Di Pardo et al., 2013). The M1-M2 polarization is partially mediated by nuclear factor kappa-light-chain-enhancer of activated B-cells (NF-κB) (Tugal et al., 2013). Activation of the NF-κB p65 subunit is critical for development of M1-associated functions in macrophages and NF-κB p65 is increased in HD patients before clinical (Di Pardo et al., 2013). Thus, one putative mechanism of aberrant pro-inflammatory responses observed during HD is the dysregulation of NF-κB activation in monocytes. The switch of monocytes to acquire a more M2-like phenotype later in the disease course could indicate a compensatory change to counteract the sustained NF-κB activation and M1-associated inflammatory state early in HD.

Compared to monocytes, less is known about how mHtt impacts the inflammatory status and the responsiveness of microglial cells to immune stimulation. Mouse models of HD demonstrate that mHtt can modify microglial responses. Lipopolysaccharide (LPS) treatment of YAC128 HD mice, which express full-length mHtt, have increased microglial cell activation compared to wild-type litter-mates (Franciosi et al., 2012). N171-82Q HD mice have increased brain indoleamine-2,3-dioxygenase, a microglial enzyme activated by inflammation, and an altered response to the protozoan *Toxoplasma gondii* (Donley et al., 2016). Therefore, based on previous studies, it is our perspective that mHtt presence is sufficient to activate monocytes/microglia and alter both the inflammatory profile and responses to extrinsic immune stimulation.

## NF-κB pathway activation in Huntington’s disease

Mutant huntingtin alters inflammatory responses in part through the NF-κB pathway as demonstrated by decreased serum IL-6 in HD mice after NF-κB suppression (Garcia-Miralles et al., 2016). It binds IKKγ, leading to increased IκB degradation and increased NF-κB signaling resulting in increased pro-inflammatory cytokine levels, including increased IL-6 (Träger et al., 2014). Mutant huntingtin also triggers mislocalization of NF-κB in synapses of neurons (Marcora and Kennedy, 2010). Decreasing NF-κB nuclear translocation is protective in HD mice further demonstrating its importance (Garcia-Miralles et al., 2016, Marcora and Kennedy, 2010). NF-κB signaling pathway can be triggered by pattern-recognition receptors (PRR) on innate immune cells that recognize pathogen-associated molecular patterns (PAMP) (Newton and Dixit, 2012, An et al., 2002). These PRR include the toll-like receptors (TLRs). LPS is an immunogenic PAMP of gram-negative bacteria such as *E. coli* that is recognized by toll-like receptor 4 (TLR4) resulting in NF-κB signaling and increased production of inflammatory mediators including IL-6 and inducible nitric oxide synthase (iNOS) (Chanput et al., 2010, Thirunavukkarasu et al., 2006, Libermann and Baltimore, 1990, An et al., 2002, Chow et al., 1999). Signaling from IL-6 (and other cytokines) then results in upregulation of iNOS via JAK/STAT (IL-6 specifically via STAT3) signaling, leading to elevated nitric oxide (NO) production (Dawn et al., 2004, Yu et al., 2003). Interleukin-6 can also stimulate NF-κB independent of LPS (Wang et al., 2003). Whether IL-6 directly upregulates NF-κB through JAK/STAT family transcription factor or through an indirect mechanism such as regulation of IKK is not clear (Lee et al., 2009, Wang et al., 2003, Yang et al., 2007). However, the evidence suggests that STAT3 synergizes with NF-κB resulting in increased pro-inflammatory cytokines including TNF-α, IL-1, and IL-6 that are increased in HD (Yang et al., 2007, Grivennikov and Karin, 2010, Björkqvist et al., 2008).

These findings support the possibility that if mHtt increases activation of NF-κB then synergy with other pathways, such as STAT signaling, could promote inflammation in HD. Evidence suggests that STAT-dependent cytokine signaling may synergize with NF-κB-activating PRR signaling to produce a hyper-responsive state in HD monocytes and microglia treated with LPS (NF-κB-activating) and IFN-γ (STAT-activating) (Björkqvist et al., 2008). However, it is unknown which signaling mechanism(s), PRR or JAK/STAT, are altered by mHtt and whether mHtt differentially impacts these pathways. Therefore, here we tested whether mHtt expressing microglial cells have altered responses to LPS to model PRR stimulation, and also to IL-6 stimulation to model STAT-mediated immune stimulation.

## Mutant huntingtin expression alters microglial responses to immune stimuli

The mouse microglial cell line EOC-20 (American Type Culture Company, CTRL-2469) was utilized as they have previously been used to study responses to inflammatory stimulus (Hensley et al., 2003, Mencel et al., 2013, Guadagno et al., 2013, Walker et al., 1995). Cells were cultured at 37°C and 5% CO_2_ in high glucose DMEM media supplemented with 20% v/v LADMAC-conditioned media according to ATCC guidelines. LADMAC cells are bone marrow-derived cells that produce high amounts of colony stimulating factor-1 needed to support EOC-20 cell growth (Olivas et al., 1995). Conditioned media was collected and frozen at −20°C until use. Wild-type (N181-18Q) or mHtt (N171-82Q) protein, or eGFP were expressed in E0C-20 cells using independent lentiviruses that use the phosphoglycerate kinase promoter. Huntingtin constructs expressed the first 3 introns of human HTT, including the CAG tract in exon 1. For each of the three viruses, a four-plasmid system was used; plasmids were transfected into HEK293T cells, virus was harvested from the media, quantified, and stored as previously described (Fox et al., 2015).

We evaluated viral transduction efficiency using eGFP-encoding virus. We identified 77.8% ± 2.5 (means ± SE, n=4) GFP-positive cells 24 hours post transduction (**Fig. 1A**). We further verified N171-18/82Q expression using the MAB5492 (EMD Millipore) (**Figs. 1B-C**). Inclusions were not observed. Because the antibody also identifies endogenous murine huntingtin we quantified total cellular fluorescence on a per-cell basis in mock transduced and N171 huntingtin expressing cells. N171-18/82Q expression resulted in a greater than 2-fold increase in expression over levels attributed to endogenous huntingtin protein (**Fig. 1D**). There was no effect of N171-18/82Q expression on cell viability compared to GFP as measured using a lactate dehydrogenase release assay (**Fig. 1E**). The microglial culture model therefore provides a way to assess the effects of mHtt on intrinsic responses to inflammatory stimuli.

**Figure 1.**
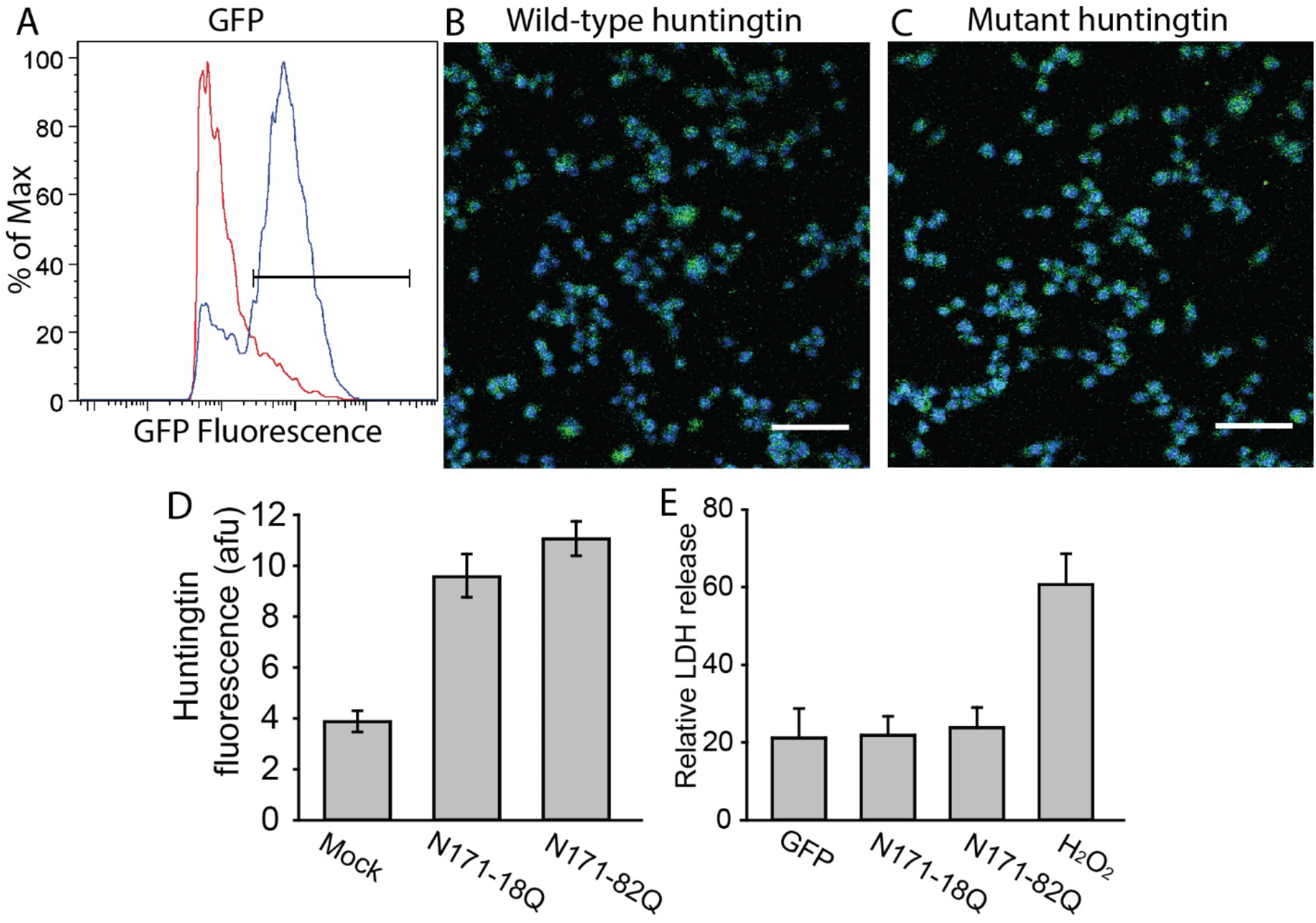
Lentiviral-mediated huntingtin expression in cultured microglial cells. EOC-20 cells were transduced for 24 hours by adding 10 ng/ml p24 viral equivalents to the media with 8 μg/ml hexadimethrine bromide (Sigma). **A.** Transduction efficiency as determined by the proportion of GFP-expressing cells, quantified using a Guava easyCyte 12HT flow cytometer. After transduction in 6-well plates, cells were scraped in PBS, counted, and plated for analysis at 5.0×10^5^ cells/well. Dead cells were labeled with Fixable Near-IR Live/Dead stain (Life Technologies) and gated from the final analysis. Representative fluorescence histogram shows the gating strategy for identifying GFP expressing cells. Red = mock transduced cells; blue = eGFP transduced cells. **B-C.** Confocal photomicrographs showing N171-18Q (B) and N171-82Q (C) huntingtin expression in EOC-20 cells. Blue = dapi, green = huntingtin; scale bars = 50 μm. **D.** Fluorescence was quantified on a per-cell basis using DAPI to identify cells from immunofluorescence images. N=152 cells from three mock transduced wells, n=170 cells from three N171-18Q transduced wells, and n=199 cells from three N171-82Q transduced wells. Bars represent means ± 95% CI. **E.** Expression of N171-82Q does not alter cell viability as determined by LDH release. Lactate dehydrogenase (LDH) activity was measured in the cell fraction and medium. Relative LDH release was determined as the activity in the supernatant divided by the total activity in supernatant and cell fraction. Two percent hydrogen peroxide was used as a positive control. N=7 for GFP, N171-18Q, N171-82Q groups and N=4 for H_2_O_2_ control. Bars represent means ± standard errors.

Effects of wild-type huntingtin and mHtt on the microglial cells was assessed by measuring responses to *E. coli* LPS (Sigma) and interleukin-6 (Biolegend). We measured nitrite, iNOS and NF-κB levels in response to these; additionally we measured IL-6 production after LPS stimulation. Nitrite is an oxidative product of NO that is further oxidized in medium to nitrate. Nitrates were reduced to nitrite using *Aspergillus* nitrate reductase and the Griess test was used to quantify nitrite (Guevara et al., 1998, Gilliam et al., 1993). Total nitrite/nitrate (hereafter called nitrite) is a surrogate for iNOS activity. Standards were made using sodium nitrite in cell culture media. IL-6 was quantified using an ELISA according to manufacturer’s guidelines (Biolegend). Relative iNOS was quantified using an ELISA (MyBioSource) (Mendonca et al., 2017). Activation of NF-κB was quantified using an ELISA for total and phosphorylated p65 subunit levels (Thermo Fisher) (Roth-Walter et al., 2014). The ratio of phosphorylated to total NFκB p65 is presented with increasing ratio indicative of activation. As expected both GFP and N171-18Q expressing cells demonstrated significantly increased IL-6 and nitrite levels in response to LPS. However, N171-82Q expressing cells lacked a response to LPS (**Figs. 2A-B**). Consistent with decreased nitrite levels following LPS stimulation, iNOS levels in N171-82Q cells did not increase following LPS treatment (**Fig. 2C**). Since LPS activates NF-κB which then upregulates expression of IL-6 and iNOS we quantified NF-κB activation in our cells with and without LPS stimulation. Baseline NF-κB activation measured by p65 subunit phosphorylation was increased in N171-82Q expressing cells compared to GFP and N171-18Q expressing cells (**Fig. 2D**). Low dose LPS increased NF-κB in GFP and N171-18Q expressing cells only while high does LPS increased NF-κB in all groups compared to no LPS controls (**Fig. 2D**).

**Figure 2.**
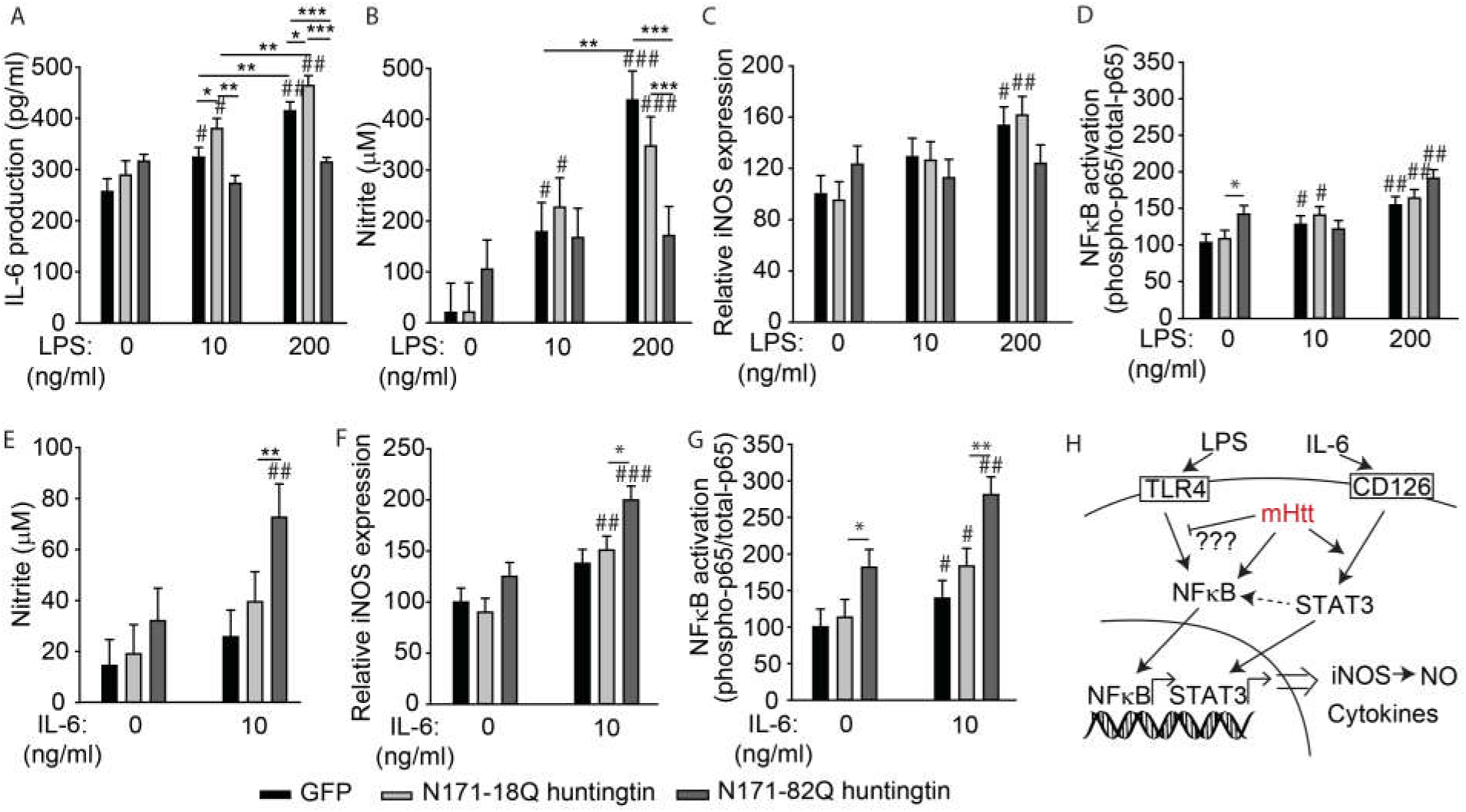
Mutant huntingtin alters the response of cultured microglial cells to immune stimulation. Culture media was changed 24 hours after transduction then BOG-20 cells were stimulated with LPS or IL-6 for 12 hours (**A-C, E-F**) or 2 hours (**D, G**) before analysis. Cells were lysed then IL-6 and nitrite measured in the supernatant fractions. Experimental values for each well were determined from technical duplicates. n=independent samples. Bars: left to right; black=GFP, light gray=N171-18Q, and, dark gray=N171-82Q. **A-B.** GFP and N171-18Q expressing cells dose-dependently respond to LPS stimulation by increasing IL-6 and nitrite production. In contrast, N171-82Q-expressing cells fail to upregulate IL-6 in response to LPS (**A,** n=6) and have less nitrite production (**B,** n=8). **C.** GFP and wild-type huntingtin-expressing, but not mHtt-expressing cells upregulate iNOS in response to LPS (n=5). **D.** Cells expressing mHtt have higher NF-κB activation at baseline and altered responses to LPS compared to wild-type expressing cells (n=6). **E.** Cells expressing mHtt have increased nitrite production after IL-6 stimulation (n=6). **F.** Mutant huntingtin-expressing cells have increased iNOS in response to IL-6 treatment compared to wild-type huntingtin-expressing cells. **G.** Cells expressing mHtt have higher NF-κB activation at baseline and with IL-6 stimulation compared to wild-type expressing cells (n=6). Bars=means ± SE. P-values: #<0.05, ##<0.01, and ###<0.001 comparison to no LPS or IL6 treatment within the same group. *=p<0.05, **=p<0.01, ***=p<0.001. Analyses were performed using the GLM procedure in SAS. Both iNOS and NF-κB relative levels were normalized to GFP-transduced cells. **H.** Model figure. Mutant huntingtin alters inflammatory signaling in microglia resulting in decreased responses to LPS and increased response to IL-6 stimulation.

We then tested the effect of STAT pathway-inducing IL-6 on iNOS and NF-κB activation. IL-6 did not have a significant effect in GFP and N171-18Q groups; however, N171-82Q cells demonstrated significantly increased nitrite production and iNOS levels (**Figs. 2E-F**). N171-82Q expressing cells also demonstrated increased NF-κB activation after IL-6 stimulation (**Fig. 2G**) and had a greater level of NF-κB activation compared to low dose LPS (**Fig. 2D**). These results indicate that mHtt sensitizes these cells to IL-6 stimulation, an effect opposite to that of LPS (see model **Fig. 2H**).

## Mutant Huntingtin impacts microglial immune signaling

Mutant huntingtin promotes increased inflammation and microglial cell activation in human HD (Crotti et al., 2014, Björkqvist et al., 2008, Tai et al., 2007). However, the ways that mHtt drives this process, especially in microglial cells is unclear. Mutant huntingtin could affect microglial cell inflammatory processes in the absence of immune stimulation, and/or alter their responses to immune stimulants. Here we assessed the effect of mHtt on cultured microglial cell functional responses to LPS and IL-6-mediated stimulation. Not only did mHtt alter microglial immune responses, but the direction of the response depended on the nature of stimulation (**Fig. 2**).

LPS signals TLR4-mediated NF-κB activation results in induction of iNOS and IL-6 (Libermann and Baltimore, 1990, Arias-Salvatierra et al., 2011). The low dose of LPS we used is established to activate NF-κB (Sung et al., 2014). The lack of increased NF-κB activation with low dose LPS stimulation in N171-82Q expressing cells together with decreased nitrite and IL-6 suggests that mHtt induces tolerance similar to LPS-induced models of tolerance in macrophages. LPS-induced macrophage tolerance is characterized, among other mechanisms, by increases in the suppressive p50 NF-κB subunit that reduces inflammation by blocking NF-κB-p65 (Kastenbauer and Ziegler-Heitbrock, 1999, Rajaiah et al., 2013). Therefore, while NF-κB-p65 is increased, cells are “tolerized” as displayed by decreased functionality which can be overcome by high doses of LPS stimulation.

HD monocytes and microglia have increased IL-6 production when stimulated with both LPS and IFN-γ, a STAT activator (Björkqvist et al., 2008). In the studied microglial cell line, mHtt had disparate effects on these responses (**Fig. 2H**). We observed that mHtt rendered the microglial cells hyporesponsive to low-dose LPS, but hypersensitive to IL-6, a cytokine that activates STAT pathways (**Figs. 2E-H**). STAT3 signaling downstream of IL-6 can synergize with NF-κB to promote inflammation (Grivennikov and Karin, 2010, Yang et al., 2007). Therefore, one possibility is that mHtt, despite reducing PRR signaling downstream of LPS stimulation, could promote synergy of NF-κB (from a PRR agonist) and STAT transcription factors (from cytokine signaling) and enhance immune activation. Our data suggests the possibility of a novel mechanism underlying HD-associated neuroinflammation driven by interaction of NF-κB and STAT signaling pathways and/or the differential effect of mHtt on these pathways.

The findings also suggest a possible mechanism where IL-6 downstream of LPS stimulation is necessary in microglial cells to drive iNOS activity and NO production. When mHtt-expressing cells are stimulated with IL-6, NF-κB activation and nitrite are greatly increased. Given these data, we think that the IL-6 requirement for iNOS activity in microglia is not a complete explanation of our results. Therefore our perspective, taking into consideration previously published studies, is that mHtt has disparate effects on PRR signaling and STAT signaling pathways that differentially impacts sensitivity of cells to extrinsic stimuli which impacts microglial activation. Cell culture studies may not replicate all aspects of endogenous microglia, therefore additional studies are needed to determine if similar effects of mHtt on microglia occur in other microglial culture models and *in vivo*.

## Conclusion

HD is only caused by a mutation within the *HTT* gene. However, there is significant variability in age of onset, after adjusting for the size of the CAG repeat size, that is partly explained by environmental factors (USVCRP and Wexler, 2004). There is considerable evidence that mHtt results in increased baseline inflammation (Björkqvist et al., 2008, Träger et al., 2014, Crotti et al., 2014). However, recent evidence also points towards altered responses to a common infection in a mouse model of HD (Donley et al., 2016). Here we demonstrate disparate effects of LPS and IL-6 on microglial responses in mHtt expressing microglial cells. These finding suggest that mHtt may also alter responses to other immune molecules, microbial antigens, or neurotropic pathogens. If these mechanisms apply to human HD then these could contribute significantly to modifying age of onset or progression by altering systemic and brain inflammatory pathways. More work is clearly needed to fully understand the mechanisms underlying altered immune responses in HD cells. However, the findings contribute to understanding mechanisms of mHtt-induced neuroinflammation and neurodegeneration as well as the variability in human HD.

## Acknowledgements

Funding was provided by University of Wyoming Neuroscience Center Grant (P30-GM103398), National Institutes of Health grant (R56-NS097813), and funds internal to the Department of Veterinary Sciences. The Wyoming INBRE 2 award (NCRR-P20RR016474/ NIGMS-P20GM103432) provided an undergraduate research fellowship to RN and a graduate student fellowship to DWD. We thank Dr. Donald Jarvis for use of laboratory equipment.

## References

An, H., Yu, Y., Zhang, M., Xu, H., Qi, R., Yan, X., Liu, S., Wang, W., Guo, Z., Guo, J., Qin, Z. & Cao, X. 2002. Involvement of ERK, p38 and NF-κB signal transduction in regulation of TLR2, TLR4 and TLR9 gene expression induced by lipopolysaccharide in mouse dendritic cells. Immunology, 106, 38–45.

Arias-Salvatierra, D., Silbergeld, E.K., Acosta-Saavedra, L.C. & Calderon-Aranda, E. S. 2011. Role of nitric oxide produced by iNOS through NF-kappaB pathway in migration of cerebellar granule neurons induced by Lipopolysaccharide. Cell Signal, 23, 425–35.

BjöRkqvist, M., Wild, E.J., Thiele, J., Silvestroni, A., Andre, R., Lahiri, N., Raibon, E., Lee, R.V., Benn, C.L., Soulet, D., Magnusson, A., Woodman, B., Landles, C., Pouladi, M.A., Hayden, M.R., Khalili-Shirazi, A., Lowdell, M.W., Brundin, P., Bates, G.P., Leavitt, B.R., MöLler, T. & Tabrizi, S. J. 2008. A novel pathogenic pathway of immune activation detectable before clinical onset in Huntington’s disease. The Journal of Experimental Medicine, 205, 1869–1877.

Chanput, W., Mes, J., Vreeburg, R.A., Savelkoul, H.F. & Wichers, H. J. 2010. Transcription profiles of LPS-stimulated THP-1 monocytes and macrophages: a tool to study inflammation modulating effects of food-derived compounds. Food Funct, 1, 254–61.

Chow, J. C., Young, D.W., Golenbock, D.T., Christ, W.J. & Gusovsky, F. 1999. Toll-like Receptor-4 Mediates Lipopolysaccharide-induced Signal Transduction. Journal of Biological Chemistry, 274, 10689–10692.

Cisbani, G. & Cicchetti, F. 2012. An in vitro perspective on the molecular mechanisms underlying mutant huntingtin protein toxicity. Cell Death Dis, 3, e382.

Crotti, A., Benner, C., Kerman, B.E., Gosselin, D., Lagier-Tourenne, C., Zuccato, C., Cattaneo, E., Gage, F.H., Cleveland, D.W. & Glass, C. K. 2014. Mutant Huntingtin promotes autonomous microglia activation via myeloid lineage-determining factors. Nat Neurosci, 17, 513–521.

Dawn, B., Xuan, Y.-T., Guo, Y., Rezazadeh, A., Stein, A.B., Hunt, G., Wu, W.-J., Tan, W. & Bolli, R. 2004. IL-6 plays an obligatory role in late preconditioning via JAK-STAT signaling and upregulation of iNOS and COX-2. Cardiovascular Research, 64, 61–71.

Di Pardo, A., Alberti, S., Maglione, V., Amico, E., Cortes, E.P., Elifani, F., Battaglia, G., Busceti, C.L., Nicoletti, F., Vonsattel, J.P. G. & Squitieri, F. 2013. Changes of peripheral TGF-β1 depend on monocytes-derived macrophages in Huntington disease. Molecular Brain, 6, 55–55.

Donley, D. W., Olson, A.R., Raisbeck, M.F., Fox, J.H. & Gigley, J. P. 2016. Huntingtons Disease Mice Infected with Toxoplasma gondii Demonstrate Early Kynurenine Pathway Activation, Altered CD8+ T-Cell Responses, and Premature Mortality. PLoS One, 11, e0162404.

Fox, J., Lu, Z. & Barrows, L. 2015. Thiol-disulfide Oxidoreductases TRX1 and TMX3 Decrease Neuronal Atrophy in a Lentiviral Mouse Model of Huntington’s Disease. PLoS Currents, 7, ecurrents.hd.b966ec2eca8e2d89d2bb4d020be4351e.

Franciosi, S., Ryu, J.K., Shim, Y., Hill, A., Connolly, C., Hayden, M.R., Mclarnon, J.G. & Leavitt, B. R. 2012. Age-dependent neurovascular abnormalities and altered microglial morphology in the YAC128 mouse model of Huntington disease. Neurobiology of Disease, 45, 438–449.

Garcia-Miralles, M., Hong, X., Tan, L.J., Caron, N.S., Huang, Y., To, X.V., Lin, R.Y., Franciosi, S., Papapetropoulos, S., Hayardeny, L., Hayden, M.R., Chuang, K.-H. & Pouladi, M. A. 2016. Laquinimod rescues striatal, cortical and white matter pathology and results in modest behavioural improvements in the YAC128 model of Huntington disease. Scientific Reports, 6, 31652.

Gilliam, M. B., Sherman, M.P., Griscavage, J.M. & Ignarro, L. J. 1993. A Spectrophotometric Assay for Nitrate Using NADPH Oxidation by Aspergillus Nitrate Reductase. Analytical Biochemistry, 212, 359–365.

Greter, M., Lelios, I. & Croxford, A. L. 2015. Microglia Versus Myeloid Cell Nomenclature during Brain Inflammation. Frontiers in Immunology, 6, 249.

Grivennikov, S. & Karin, M. 2010. Dangerous liaisons: STAT3 and NF-κB collaboration and crosstalk in cancer. Cytokine & growth factor reviews, 21, 11–19.

Guadagno, J., Xu, X., Karajgikar, M., Brown, A. & Cregan, S. P. 2013. Microglia-derived TNF[alpha] induces apoptosis in neural precursor cells via transcriptional activation of the Bcl-2 family member Puma. Cell Death Dis, 4, e538.

Guevara, I., Iwanejko, J., DembiŃSka-KieĆ, A., Pankiewicz, J., Wanat, A., Anna, P., GoŁBek, I., BartuŚ, S., Malczewska-Malec, M. & Szczudlik, A. 1998. Determination of nitrite/nitrate in human biological material by the simple Griess reaction. Clinica Chimica Acta, 274, 177–188.

Hensley, K., Fedynyshyn, J., Ferrell, S., Floyd, R.A., Gordon, B., Grammas, P., Hamdheydari, L., Mhatre, M., Mou, S., Pye, Q.N., Stewart, C., West, M., West, S. & Williamson, K. S. 2003. Message and protein-level elevation of tumor necrosis factor α (TNFα) and TNFα-modulating cytokines in spinal cords of the G93A-SOD1 mouse model for amyotrophic lateral sclerosis. Neurobiology of Disease, 14, 74–80.

Kastenbauer, S. & Ziegler-Heitbrock, H. W. 1999. NF-kappaB1 (p50) is upregulated in lipopolysaccharide tolerance and can block tumor necrosis factor gene expression. Infect Immun, 67, 1553–9.

Kraft, A. D., Kaltenbach, L.S., Lo, D.C. & Harry, G. J. 2012. Activated microglia proliferate at neurites of mutant huntingtin-expressing neurons. Neurobiology of Aging, 33, 621.e17–621.e33.

Kwan, W., TrÄGer, U., Davalos, D., Chou, A., Bouchard, J., Andre, R., Miller, A., Weiss, A., Giorgini, F., Cheah, C., MÖLler, T., Stella, N., Akassoglou, K., Tabrizi, S.J. & Muchowski, P. J. 2012. Mutant huntingtin impairs immune cell migration in Huntington disease. The Journal of Clinical Investigation, 122, 4737–4747.

Lee, H., Herrmann, A., Deng, J., Kujawski, M., Niu, G., Li, Z., Forman, S., Jove, R., Pardoll, D. & Yu, H. 2009. Persistently-activated Stat3 maintains constitutive NF-κB activity in tumors. Cancer cell, 15, 283–293.

Libermann, T. A. & Baltimore, D. 1990. Activation of interleukin-6 gene expression through the NF-kappa B transcription factor. Molecular and Cellular Biology, 10, 2327–2334.

Marcora, E. & Kennedy, M. B. 2010. The Huntington’s disease mutation impairs Huntingtin’s role in the transport of NF-κB from the synapse to the nucleus. Human Molecular Genetics, 19, 4373–4384.

Mencel, M., Nash, M. & Jacobson, C. 2013. Neuregulin Upregulates Microglial α7 Nicotinic Acetylcholine Receptor Expression in Immortalized Cell Lines: Implications for Regulating Neuroinflammation. PLOS ONE, 8, e70338.

Mendonca, P., Taka, E., Bauer, D., Cobourne-Duval, M. & Soliman, K. F. A. 2017. The attenuating effects of 1,2,3,4,6 penta-O-galloyl-β-<span class=“small”>d</span>-glucose on inflammatory cytokines release from activated BV-2 microglial cells. Journal of Neuroimmunology, 305, 9–15.

Mills, C. D., Kincaid, K., Alt, J.M., Heilman, M.J. & Hill, A. M. 2000. M-1/M-2 Macrophages and the Th1/Th2 Paradigm. The Journal of Immunology, 164, 6166–6173.

Newton, K. & Dixit, V. M. 2012. Signaling in Innate Immunity and Inflammation. Cold Spring Harbor Perspectives in Biology, 4, a006049.

Olivas, E., Chen, B.B.D.M. & Walker, W. S. 1995. Use of the Pannell-Milstein roller bottle apparatus to produce high concentrations of the CSF-1, the mouse macrophage growth factor. Journal of Immunological Methods, 182, 73–79.

Rajaiah, R., Perkins, D.J., Polumuri, S.K., Zhao, A., Keegan, A.D. & Vogel, S. N. 2013. Dissociation of Endotoxin Tolerance and Differentiation of Alternatively Activated Macrophages(). Journal of immunology (Baltimore, Md. : 1950), 190, 4763–4772.

Roth-Walter, F., Moskovskich, A., Gomez-Casado, C., Diaz-Perales, A., Oida, K., Singer, J., Kinaciyan, T., Fuchs, H.C. & Jensen-Jarolim, E. 2014. Immune Suppressive Effect of Cinnamaldehyde Due to Inhibition of Proliferation and Induction of Apoptosis in Immune Cells: Implications in Cancer. PLOS ONE, 9, e108402.

Sapp, E., Kegel, K.B., Aronin, N., Hashikawa, T., Uchiyama, Y., Tohyama, K., Bhide, P.G., Vonsattel, J. P. & Difiglia, M. 2001. Early and Progressive Accumulation of Reactive Microglia in the Huntington Disease Brain. Journal of Neuropathology & Experimental Neurology, 60, 161–172.

Sung, M.-H., Li, N., Lao, Q., Gottschalk, R.A., Hager, G.L. & Fraser, I. D. C. 2014. Switching of the Relative Dominance between Feedback Mechanisms in Lipopolysaccharide-Induced NF-κB Signaling. Science signaling, 7, ra6–ra6.

Tai, Y. F., Pavese, N., Gerhard, A., Tabrizi, S.J., Barker, R.A., Brooks, D.J. & Piccini, P. 2007. Microglial activation in presymptomatic Huntington’s disease gene carriers. Brain, 130, 1759–1766.

Thirunavukkarasu, C., Watkins, S.C. & Gandhi, C. R. 2006. Mechanisms of endotoxin-induced NO, IL-6, and TNF-α production in activated rat hepatic stellate cells: Role of p38 MAPK. Hepatology, 44, 389–398.

TräGer, U., Andre, R., Lahiri, N., Magnusson-Lind, A., Weiss, A., Grueninger, S., Mckinnon, C., Sirinathsinghji, E., Kahlon, S., Pfister, E.L., Moser, R., Hummerich, H., Antoniou, M., Bates, G.P., Luthi-Carter, R., Lowdell, M.W., BjÖRkqvist, M., Ostroff, G.R., Aronin, N. & Tabrizi, S. J. 2014. HTT-lowering reverses Huntington’s disease immune dysfunction caused by NFκB pathway dysregulation. Brain, 137, 819–833.

Tugal, D., Liao, X. & Jain, M. K. 2013. Transcriptional Control of Macrophage Polarization. Arteriosclerosis, Thrombosis, and Vascular Biology, 33, 1135–1144.

Usvcrp, T. U. S.-V. C. R. P. & Wexler, N. S. 2004. Venezuelan kindreds reveal that genetic and environmental factors modulate Huntington’s disease age of onset. Proceedings of the National Academy of Sciences of the United States of America, 101, 3498–3503.

Walker, W. S., Gatewood, J., Olivas, E., Askew, D. & Havenith, C. E. G. 1995. Mouse microglial cell lines differing in constitutive and interferon-γ-inducible antigen-presenting activities for naive and memory CD4+ and CD8+ T cells. Journal of Neuroimmunology, 63, 163–174.

Wang, L., Walia, B., Evans, J., Gewirtz, A.T., Merlin, D. & Sitaraman, S. V. 2003. IL-6 Induces NF-κB Activation in the Intestinal Epithelia. The Journal of Immunology, 171, 3194–3201.

Weiss, A., TrÄGer, U., Wild, E.J., Grueninger, S., Farmer, R., Landles, C., Scahill, R.I., Lahiri, N., Haider, S., Macdonald, D., Frost, C., Bates, G.P., Bilbe, G., Kuhn, R., Andre, R. & Tabrizi, S. J. 2012. Mutant huntingtin fragmentation in immune cells tracks Huntington’s disease progression. The Journal of Clinical Investigation, 122, 3731–3736.

Yang, J., Liao, X., Agarwal, M.K., Barnes, L., Auron, P.E. & Stark, G. R. 2007. Unphosphorylated STAT3 accumulates in response to IL-6 and activates transcription by binding to NFkappaB. Genes Dev, 21, 1396–408.

Yu, X., Kennedy, R.H. & Liu, S. J. 2003. JAK2/STAT3, not ERK1/2, mediates interleukin-6-induced activation of inducible nitric-oxide synthase and decrease in contractility of adult ventricular myocytes. J Biol Chem, 278, 16304–9.

